# An antibody-escape calculator for mutations to the SARS-CoV-2 receptor-binding domain

**DOI:** 10.1101/2021.12.04.471236

**Authors:** Allison J. Greaney, Tyler N. Starr, Jesse D. Bloom

## Abstract

A key goal of SARS-CoV-2 surveillance is to rapidly identify viral variants with mutations that reduce neutralization by polyclonal antibodies elicited by vaccination or infection. Unfortunately, direct experimental characterization of new viral variants lags their sequence-based identification. Here we help address this challenge by aggregating deep mutational scanning data into an “escape calculator” that estimates the antigenic effects of arbitrary combinations of mutations to the virus’s spike receptor-binding domain (RBD). The calculator can be used to intuitively visualize how mutations impact polyclonal antibody recognition, and score the expected antigenic effect of combinations of mutations. These scores correlate with neutralization assays performed on SARS-CoV-2 variants, and emphasize the ominous antigenic properties of the recently described Omicron variant. An interactive version of the calculator is at https://jbloomlab.github.io/SARS2_RBD_Ab_escape_maps/escape-calc/, and we provide a Python module for batch processing.

---

Human coronaviruses undergo antigenic evolution that erodes antibody-based neutralization (Eguia *et al*. 2021; Kistler and Bedford 2021). This antigenic evolution is already apparent for SARS-CoV-2, as new viral variants with reduced antibody neutralization have emerged only ∼2 years since the virus first started to spread in humans. A tremendous amount of experimental effort has been expended to characterize these SARS-CoV-2 variants in neutralization assays (Wang *et al*. 2021; Uriu *et al*. 2021; Lucas *et al*. 2021). Unfortunately, the rate at which new variants arise outstrips the speed at which these experiments can be performed.

A partial solution is to use deep mutational scanning experiments to *prospectively* measure how viral mutations impact antibody binding or neutralization. Deep mutational scanning can systematically measure the antigenic impacts of all possible amino-acid mutations in key regions of spike on monoclonal antibodies (Starr *et al*. 2021b; Greaney *et al*. 2021d) or sera (Greaney *et al*. 2021a). However, SARS-CoV-2 variants of concern typically have multiple mutations, and it is not feasible to experimentally characterize all combinations of mutations even via high-throughput approaches such as deep mutational scanning.

Here we take a step towards addressing this challenge by aggregating deep mutational scanning data across many antibodies to assess the impacts of mutations in the spike receptor-binding domain (RBD), which is the primary target of neutralizing antibodies to SARS-CoV-2 (Piccoli *et al*. 2020; Greaney *et al*. 2021a; Schmidt *et al*. 2021). The resulting “escape calculator” enables qualitative visualization and quantitative scoring of the antigenic effects of arbitrary combinations of mutations. Importantly, the escape calculator is based on simple transformations of direct experimental measurements, and does not involve complex black-box computational methods.

## Results

### Combining monoclonal antibody escape maps reveals correlated and independent viral antigenic mutations

A deep mutational scanning experiment can measure how all single amino-acid mutations to the SARS-CoV-2 RBD affect binding by a monoclonal antibody (Greaney *et al*. 2021d). This mutation-level information can be summarized for each RBD site, such as by taking the mean or sum of mutation-level effects at a site. Here we will work with these site-level escape maps.

As a small example to illustrate the principle behind our approach, Figure 1A shows previously reported measurements (Starr *et al*. 2021b,c) of how mutations to each RBD site affect binding by three monoclonal antibodies: LY-CoV016 (etesevimab), LY-CoV555 (bamlanivimab), and REGN10987 (imdevimab). Each antibody targets a different epitope on the RBD: LY-CoV016 targets the class 1 epitope, LY-CoV555 the class 2 epitope, and REGN10987 the class 3 epitope (Barnes *et al*. 2020; Greaney *et al*. 2021b). Because the antibodies have distinct epitopes, they are escaped by largely distinct sets of mutations: LY-CoV016 is most strongly escaped by mutations at site 417, LY-CoV555 at site 484, and REGN10987 at sites 444–446 (Figure 1A). Now imagine a polyclonal antibody mix of these three antibodies combined at equal potencies. We can generate an escape map for this hypothetical antibody mix simply by averaging the experimentally measured escape maps for the three individual antibodies, yielding the thick black line in Figure 1A. Because this polyclonal escape map is the average of the monoclonal antibody maps, its largest peaks are at the sites of strongest escape for each individual antibody: 417, 484, and 444–446.

**Figure 1.**
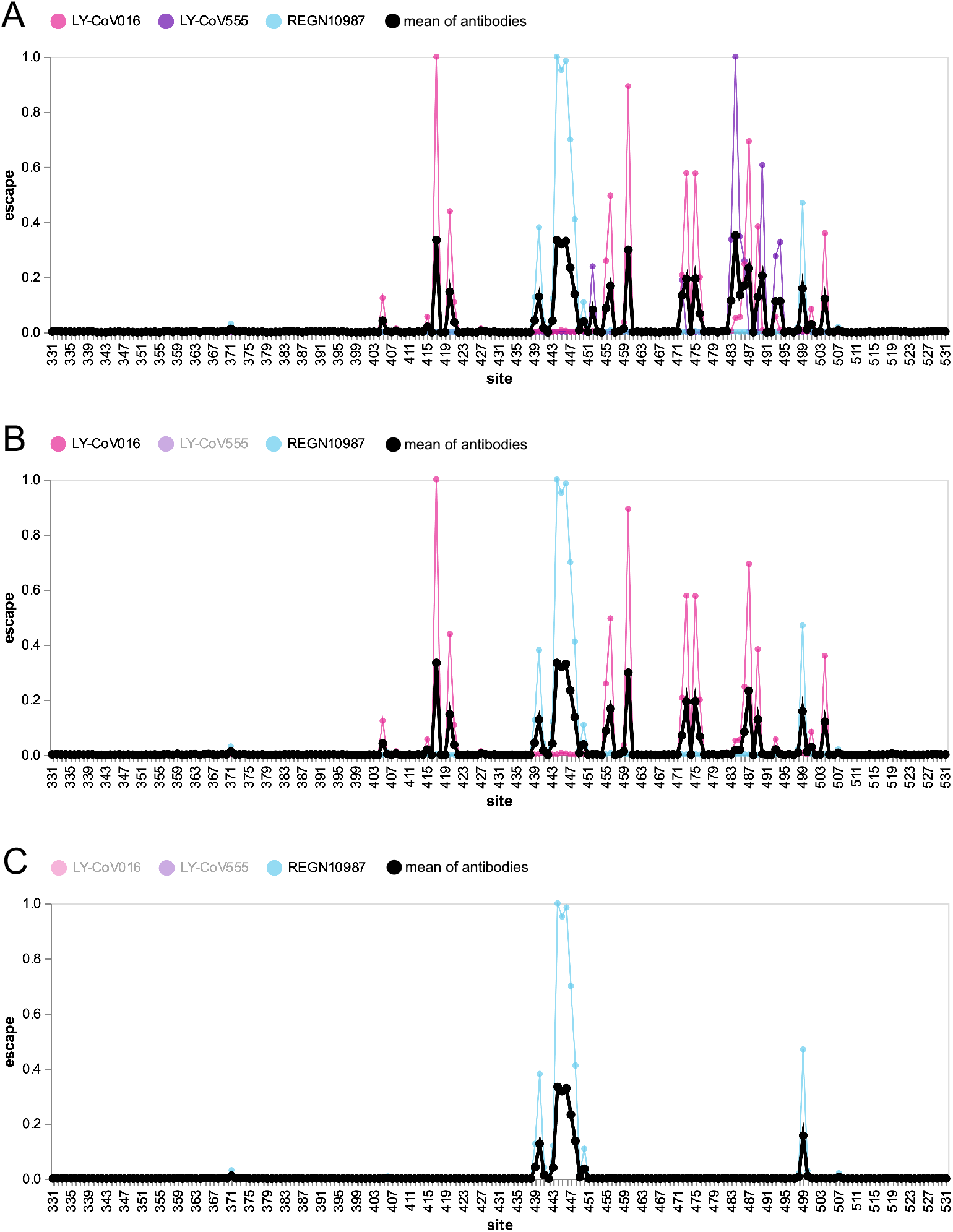
Escape map for a hypothetical polyclonal mix consisting of an equipotent mixture of three monoclonal antibodies targeting distinct epitopes on the SARS-CoV-2 RBD. **(A)** Experimentally measured escape maps for three antibodies, and the mean of these maps (thick black line). Each point on the x-axis represents a site in the RBD, and the y-axis represents the total measured escape by all mutations at that site scaled so the maximum for each antibody is one. **(B)** Escape map if the contribution of antibody LY-CoV555 is ablated.**(C)** Escape map if the contributions of antibodies LY-CoV555 and LY-CoV016 are ablated. An interactive version of this figure is at https://jbloomlab.github.io/SARS2_RBD_Ab_escape_maps/mini-example-escape-calc/.

Next imagine removing one antibody from the mix by mutating its epitope. Figure 1B shows the resulting escape map if LY-CoV555 is ablated, as would occur if site 484 was mutated. The thick black line for the antibody mix no longer has peaks at 484 and other sites targeted by LY-CoV555, such as 490. Therefore, in this hypothetical polyclonal antibody mix, escape at sites 484 and 490 is correlated since both sites are targeted by the same antibody. However, the polyclonal mix’s escape map at sites 417 and 460 is unaffected by mutations that escape LY-CoV555, since they are targeted by a different antibody, LY-CoV016. But if we also ablate LY-CoV016 (such as by mutating site 417), then the peaks at 417 and 460 also disappear, and the remaining peaks are at sites targeted by REGN10987, such as 444–446 (Figure 1C). Of course, if REGN10987 was also ablated such as by mutating site 446, then the polyclonal antibody mix would have no remaining activity. This and other scenarios can be explored using the interactive version of Figure 1 at https://jbloomlab.github.io/SARS2_RBD_Ab_escape_maps/mini-example-escape-calc/.

### Aggregating deep mutational scanning data for 33 human antibodies yields a realistic escape calculator

The toy example in the previous section illustrates how experimental data for individual antibodies can be combined to yield an escape map for a hypothetical polyclonal antibody mix. To create an escape map for an antibody mix that more realistically represents actual human sera, we aggregated previously generated deep mutational scanning data for 33 neutralizing antibodies elicited by SARS-CoV-2. These antibodies were isolated from a variety of patient cohorts within the first year of the pandemic (see Methods for details). An assumption of the analysis that follows is that an equipotent mixture of these 33 antibodies represents the neutralizing activity of human sera; we emphasize that this assumption is imperfect since in reality the antibodies were chosen for prior study for a variety of ad hoc reasons. The escape maps for all the individual antibodies can be interactively interrogated at https://jbloomlab.github.io/SARS2_RBD_Ab_escape_maps/.

The overall polyclonal escape map generated by averaging the experimental data for all 33 antibodies is in Figure 2A. As in the toy three-antibody example in the previous section, there are peaks at sites 417, 484, and 444–446. However, the peak at 484 is now larger than any other peak, reflecting the fact that antibodies targeting the class 2 epitope containing E484 are especially common in the human antibody response to early SARS-CoV-2 strains (Yuan *et al*. 2020; Robbiani *et al*. 2020; Greaney *et al*. 2021a,b; Chen *et al*. 2021). In addition, there are smaller peaks at a variety of other sites, reflecting the fact that each antibody has a somewhat idiosyncratic epitope (Figure 2A).

**Figure 2.**
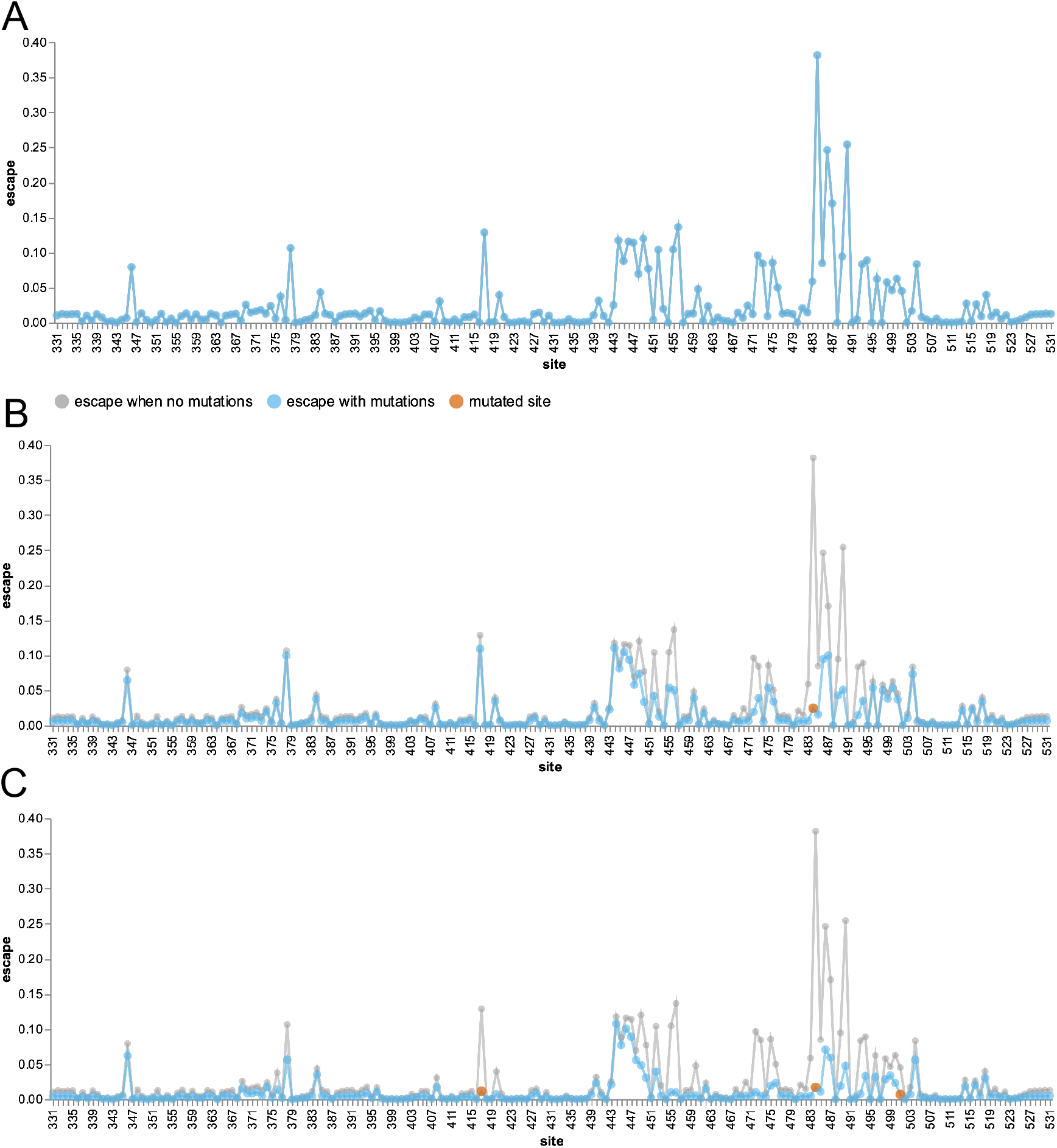
An escape calculator generated by aggregating deep mutational scanning for 33 neutralizing antibodies targeting the SARS-CoV-2 RBD. **(A)** The blue line shows the extent of escape mediated by mutations at each site, as estimated by simply averaging the data for all the individual antibodies. **(B)** The blue line shows escape map after a mutation to site 484 (red point) ablates recognition by antibodies strongly targeting that site, while the gray line shows the original escape map in the absence of any mutations. **(C)** The escape map after mutating sites 417, 484, and 501 (the three RBD sites mutated in the Beta variant). An interactive version of this figure is at https://jbloomlab.github.io/SARS2_RBD_Ab_escape_maps/escape-calc/.

We can follow the principle outlined in the toy example in the previous section to calculate the expected polyclonal escape map *after* mutating sites in the RBD. Specifically, we reduce the contribution of each antibody by an amount that scales with how strongly that antibody targets each mutated site (see Methods for details). For instance, the blue lines in Figure 2B show the polyclonal escape map after mutating site 484. Mutating site 484 obviously drops the contribution of that site, but it also decreases the contribution of other sites such as 490 that are commonly targeted by antibodies with epitopes that include site 484. In contrast, mutating site 484 has minimal effect on the polyclonal escape map at sites like 417 or 444–446, since those sites are generally targeted by antibodies that are unaffected by mutations at site 484.

We can also calculate the expected effects of compound mutations. Figure 2C shows the polyclonal escape map after mutating all three RBD sites that are changed in the Beta variant (sites 417, 484, and 501). This polyclonal escape map has lost contributions not only from the mutated sites, but also sites that form common epitopes with 417 or 484 (e.g., sites 455, 456, 486, and 490). However, the escape map still has major contributions from antibodies targeting sites like 444–446, since such antibodies are generally unaffected by mutations at sites 417, 484, or 501.

We recommend the reader explores the interactive escape calculator at https://jbloomlab.github.io/SARS2_RBD_Ab_escape_maps/escape-calc/, to perform calculations like those in Figure 2 for arbitrary combinations of mutated RBD sites. Such visual exploration of different combinations of mutations provides an intuitive sense of the antigenic structure of the RBD.

### The escape calculations correlate well with neutralization assays of human polyclonal sera against SARS-CoV-2 variants

For each set of mutated RBD sites, we can define a quantitative score that represents the polyclonal antibody binding that remains after mutating these sites. This score is defined using the same principle as the site-wise escape calculator in the previous section: we reduce the contribution of each antibody by an amount that scales with how strongly it is escaped by each mutated site, and define the overall score as the fraction of all antibody contributions that remain (see Methods for details). This calculation is implemented in the interactive calculator at https://jbloomlab.github.io/SARS2_RBD_Ab_escape_maps/escape-calc/, and returns a score that ranges from one (no mutations affect binding of any antibodies) to zero (all antibodies fully escaped).

To test how these escape-calculator scores compare to experimentally measured neutralization titers, we collated neutralization data from three previously published studies (Lucas *et al*. 2021; Uriu *et al*. 2021; Wang *et al*. 2021), each of which characterized sera from two patient cohorts against a variety of SARS-CoV-2 variants and mutants. One can imagine many reasons why the escape-calculator scores might differ from the real neutralization titers: the calculator only considers RBD mutations, the antibodies used by the calculator might not accurately reflect the real mix in polyclonal sera, etc. But despite all these potential caveats, the escape-calculator scores correlate quite well with the measured neutralization titers across all studies and cohorts (Figure 3). Therefore, the simple and intuitive approach used by the calculator seems to accurately reflect the dominant features of polyclonal antibody escape in the RBD.

**Figure 3.**
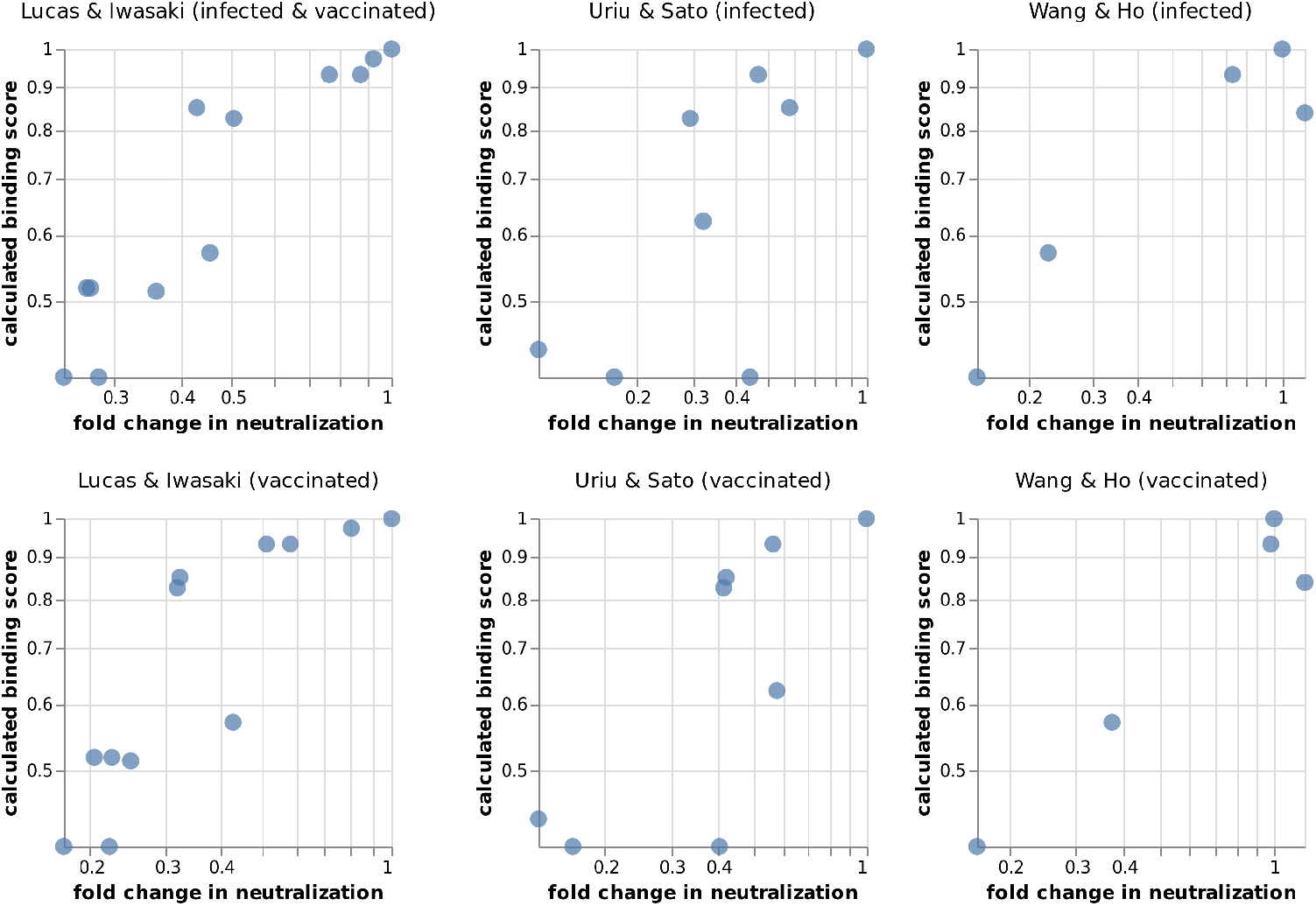
Correlation of calculated binding scores with experimentally measured fold changes in neutralization for SARS-CoV-2 variants and mutants (smaller values indicate worse neutralization). The data of Lucas *et al*. (2021) was generated using authentic SARS-CoV-2 and sera from vaccinated individuals who were (top) or were not (bottom) previously infected with SARS-CoV-2. The data of Uriu *et al*. (2021) and Wang *et al*. (2021) was generated using pseudovirus against convalescent (top) or vaccine (bottom) sera, with vaccine sera from Pfizer BNT162b2 or Moderna mRNA-1273 vaccines, respectively. The fold changes are geometric means over all subjects in each cohort. An interactive version of this figure that allows mousing over points to see details is at https://jbloomlab.github.io/RBD_escape_calculator_paper/neut_studies.html.

### The escape calculator suggests extensive antigenic change in the new Omicron variant

We applied the escape calculator to the recently reported Omicron variant, which has 15 mutated sites in its RBD (NGS-SA 2021; de Oliveira 2021). The calculated binding score for the Omicron variant is much lower than any other SARS-CoV-2 variants of concern, indicating extensive antibody escape (Figure 4A). The Omicron variant’s calculated score is roughly equivalent to that of a polymutant spike (PMS20) that was artificially engineered in a pseudovirus by Schmidt *et al*. (2021) to maximize escape from polyclonal serum antibodies. For comparison, Schmidt *et al*. (2021) measured that neutralization titers against this artificial PMS20 spike were reduced by ∼20- to ∼80-fold for sera from various cohorts of vaccinated and infected individuals.

**Figure 4.**
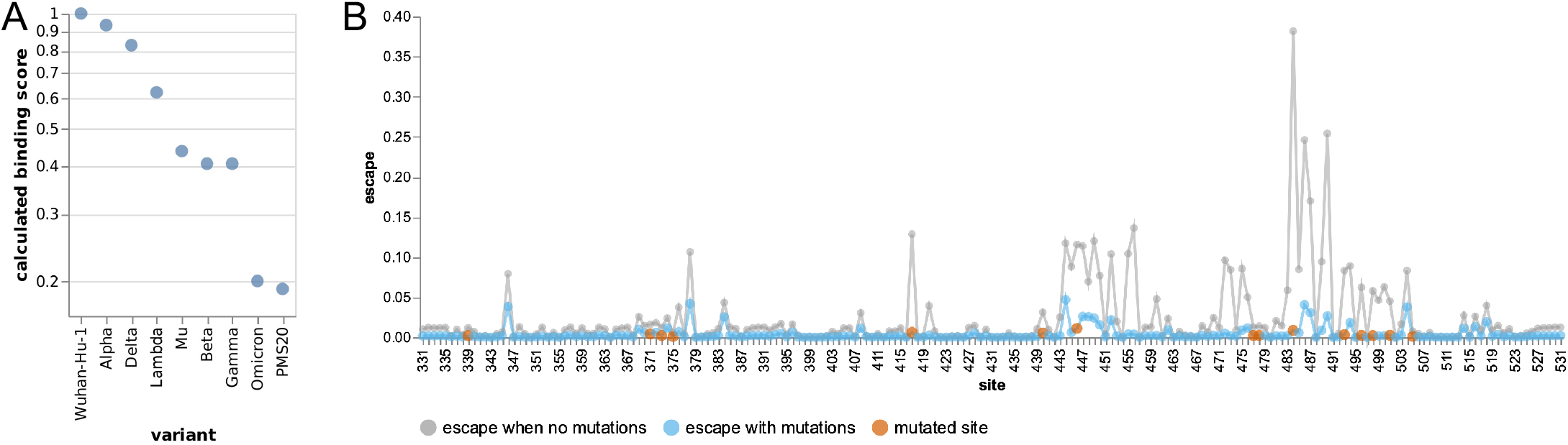
Escape calculations for the Omicron variant. **(A)** The calculated binding scores for SARS-CoV-2 variants and the artificial polymutant spike (PMS20) generated by Schmidt *et al*. (2021). Scores of one indicate no mutations affect binding, and scores of zero indicate no antibody binding remains. An interactive version of this plot that allows mousing over points to see details is at https://jbloomlab.github.io/RBD_escape_calculator_paper/variants.html. **(B)** The calculated escape map for the Omicron variant’s RBD (blue) compared to an unmutated RBD (gray), with sites of mutations in the Omicron variant in red. The mutated RBD sites for each variant are in Table 1.

The site-level escape map for the Omicron variant’s RBD is shown in Figure 4B. The Omicron RBD has lost most peaks of antibody binding relative to the original RBD. Exploration of the mutations using the interactive calculator at https://jbloomlab.github.io/SARS2_RBD_Ab_escape_maps/escape-calc/ indicates that mutations at sites 484, 446, and 417 are the biggest drivers of this antigenic change, although other mutations also contribute. The residual peaks in the map suggests the remaining antibody activity against the Omicron variant RBD could be further eroded by mutations at sites like 346, 378, 444, and 504.

## Discussion

We have described an escape calculator that uses experimental data for 33 monoclonal antibodies to estimate the antigenic effects of arbitrary combinations of mutations to the SARS-CoV-2 RBD. The key insight is to aggregate data for individual antibodies to define both which RBD sites are antigenically important, and which combinations of mutations have redundant versus additive effects on antibody binding. For instance, sites 417, 484, and 490 are all peaks of antibody escape. But mutations at 484 and 490 have redundant effects since they generally escape the same antibodies, whereas mutations at 484 and 417 have additive effects since they generally escape different antibodies.

Another key aspect of our approach is the interactive visual implementation of the escape calculator, so the user can interrogate the effects of mutations (or combinations of mutations) simply by clicking (or shift-clicking) on sites. This interactivity provides an intuitive understanding of the antigenic structure of the RBD, and shows how the calculator is performing simple transformations directly on experimental data. We encourage interactive use of the calculator so it acts as an aid to augment human interpretation, rather than a black-box computational method. However, we note that in addition to the interactive calculator at https://jbloomlab.github.io/SARS2_RBD_Ab_escape_maps/escape-calc/, we also provide a Python module for batch processing of large numbers of sequences at https://github.com/jbloomlab/SARS2_RBD_Ab_escape_maps/.

There are caveats that should be kept in mind when using the escape calculator. First, the calculator only considers sites in the RBD, and ignores mutations to other regions of spike. Second, the calculator assumes the neutralizing activity of human polyclonal serum is represented by an equipotent mix of the monoclonal antibodies that happen to have been previously characterized by deep mutational scanning. Third, the calculator simply averages site-level escape measurements across antibodies, and does not yet implement a real biophysical model of the combined activity of multiple antibodies (Einav and Bloom 2020). Finally, and in our minds most significantly, the calculator estimates the impact of mutations in reference to antibodies targeted to the early Wuhan-Hu-1 RBD—an approach that is currently reasonable, but will become problematic as human exposure and vaccination histories diversify in the years to come (see last paragraph).

Despite all these caveats, the escape calculator yields binding scores that correlate with experimentally measured neutralization titers. In addition, the actual antigenic evolution of SARS-CoV-2 seems to follow the principles captured by the escape calculator: variants of concern generally have combinations of mutations calculated to have additive effects on antibody escape (e.g., 417 and 484) rather than combinations calculated to have redundant effects (e.g., 484 and 490). We suspect the calculator works well because the RBD is the dominant target of neutralizing activity (Piccoli *et al*. 2020; Greaney *et al*. 2021a; Schmidt *et al*. 2021) and the human antibody-response to the early Wuhan-Hu-1 RBD shares broad commonalities across individuals (Yuan *et al*. 2020; Robbiani *et al*. 2020; Greaney *et al*. 2021a,b; Chen *et al*. 2021).

However, the situation will become more complex over time. Currently, most humans with antibodies to SARS-CoV-2 have been exposed to a RBD antigen that is identical or very similar to that of the early Wuhan-Hu-1 strain. Therefore, antigenic studies can reasonably define mutations in reference to that RBD, since it is what the antibodies target. But as humans are exposed to more diverged RBD variants, it will become difficult to determine what reference to use to define antigenic mutations, since different individuals will have antibodies targeting different RBDs. Additionally, differing exposure histories can leave individuals with different antibody specificities (Cobey and Hensley 2017), a process that is already starting to occur for SARS-CoV-2 (Greaney *et al*. 2021c). So in the future, it will be necessary to stratify the data used by the escape calculator by which RBD variant elicited the antibodies, and aggregate data for antibodies that reflect the sera in question. For this reason, we expect to continue adding to the data used by the escape calculator, and emphasize that it will change over time from the version described here, although we provide stable links to the current version in the Methods below.

## Methods

### Code and data availability

The most up-to-date code and data used to implement the escape calculator are at https://github.com/jbloomlab/SARS2_RBD_Ab_escape_maps, and the version described in this paper are at https://github.com/jbloomlab/SARS2_RBD_Ab_escape_maps/tree/bioRxiv_v1.

The data used by the escape calculator are at https://github.com/jbloomlab/SARS2_RBD_Ab_escape_maps/blob/main/processed_data/escape_calculator_data.csv, and the version used for this paper are at https://github.com/jbloomlab/SARS2_RBD_Ab_escape_maps/blob/bioRxiv_v1/processed_data/escape_calculator_data.csv.

A downloadable HTML version of the escape calculator is at https://github.com/jbloomlab/SARS2_RBD_Ab_escape_maps/raw/main/docs/_includes/escape_calc_chart.html, with the version described in this paper at https://github.com/jbloomlab/SARS2_RBD_Ab_escape_maps/blob/bioRxiv_v1/docs/_includes/escape_calc_chart.html.

### Interactive versions of figures

Interactive versions of all of the figures in this paper are at https://jbloomlab.github.io/RBD_escape_calculator_paper/. These figures allow mousing over points to see details, etc.

### Deep mutational scanning data used by the calculator

The experimental data used by the escape calculator are drawn from seven previously published deep mutational scanning studies (Greaney *et al*. 2021d,b; Starr *et al*. 2021b,c,a; Dong *et al*. 2021; Tortorici *et al*. 2021) and one unpublished dataset available at https://github.com/jbloomlab/SARS-CoV-2-RBD_MAP_COV2-2955. In total, these studies contain data for 36 monoclonal antibodies. Three of these antibodies (CR3022, S304, and S309) were elicited by infection with SARS-CoV-1 and so are excluded from the datasets used for the calculations in this paper, although the calculator has an option (`eliciting_virus`) that allows optional inclusion of these antibodies. The majority of the antibodies were originally isolated from cohorts of individuals infected with SARS-CoV-2 in the first half of 2020, and were initially characterized by the Crowe lab (Zost *et al*. 2020), Nussenzweig lab (Robbiani *et al*. 2020), or Vir Biotechnology (Piccoli *et al*. 2020), with a few additional antibodies coming from commercial synthesis based on previously reported sequences (Hansen *et al*. 2020; Jones *et al*. 2021; Shi *et al*. 2020). The full deep mutational scanning data for all these antibodies are interactively displayed at https://jbloomlab.github.io/SARS2_RBD_Ab_escape_maps/ and available in raw form at https://raw.githubusercontent.com/jbloomlab/SARS2_RBD_Ab_escape_maps/main/processed_data/escape_data.csv (see https://github.com/jbloomlab/SARS2_RBD_Ab_escape_maps/blob/bioRxiv_v1/processed_data/escape_data.csv for a stable version of the raw data corresponding to that used in this paper).

The deep mutational scanning measures an escape fraction for each tolerated RBD mutation against each antibody, which represents an estimate of how completely that mutation escapes antibody binding (Greaney *et al*. 2021d). We summarize the mutation-level escape fractions into site-level measurements in two ways: taking the sum of the mutation escape fractions at each site, or taking the mean of the mutation escape fractions across all tolerated mutations at each site. The results reported in this paper use the sums as the site-level metric, although the calculator has an option (`escape_metric`) to use the mean instead. We also normalize the site-level escape metrics for each antibody to account for different strengths of antibody selection in different experiments using the approach described in Greaney *et al*. (2021a) and implemented in https://jbloomlab.github.io/dmslogo/dmslogo.utils.html#dmslogo.utils.AxLimSetter with `min_upperlim=1` and `max_from_quantile=(0.5, 0.05)`: essentially this corresponds to scaling the site escape values for each antibody so that a value of one corresponds to the larger of the maximum escape at a site or 20 times the median value across sites. The results reported in this paper use the normalized site-level metrics, although the calculator has an option (`escape_values_normalized`) to also use the non-normalized values.

### Calculation of the impact of mutations

The escape calculator determines the impact of mutating sites by calculating how much each antibody is escaped by mutations at each site, and adjusting its contribution to the overall polyclonal mix accordingly. Specifically, for each antibody *a* we have a deep mutational scanning measurement *x*_*a,r*_ of how much mutating *r* escapes that antibody. In the absence of any mutations, the overall escape map shown for the polyclonal mix is simply the mean over all antibodies, 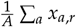 where *A* is the number of antibodies.

Let *ℳ* be the set of sites that are mutated. Then for each antibody we compute the binding retained as 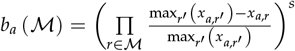. Essentially, this equation means that if the RBD is mutated at a strong site of escape for an antibody *a*, much if the binding is lost (if it is the strongest site of escape, all binding is lost). The variable *s* represents how dramatically binding is lost for mutations at sites of escape that are not the strongest: larger values of *s* means mutations at even moderate sites of escape reduce binding a lot. In this paper we report calculations with *s* = 2, although the calculator has an option (`mutation_escape_-strength`) to choose other values. We then define the escape map after the mutations *ℳ* as 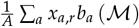. The calculator shows the escape map with no mutations in gray, and that after mutations in blue.

The overall antibody binding scores represent the fraction of antibodies that still bind, and are calculated simply as 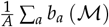.

### Implementation of interactive calculator

The interactive calculator at https://jbloomlab.github.io/SARS2_RBD_Ab_escape_maps/escape-calc/ is imlemented using Altair (https://altair-viz.github.io/) (VanderPlas *et al*. 2018), which is in turn built upon Vega-Lite (Satyanarayan *et al*. 2017).

### Python module with batch-mode calculator

A Python module that implements the calculations is at https://github.com/jbloomlab/SARS2_RBD_Ab_escape_maps/, and has all the same options as the interactive calculator.

### Compilation of neutralization titers from the literature

For Figure 3, we compiled neutralization data from three published studies on SARS-CoV-2 variants and mutants (Lucas *et al*. 2021; Uriu *et al*. 2021; Wang *et al*. 2021). For each study cohort, we computed the geometric mean fold change in neutralization titer over all subjects. The numerical compiled data are at https://github.com/jbloomlab/RBD_escape_calculator_paper/tree/main/results/neut_studies.

### Mutations in SARS-CoV-2 variants

For Figure 4, the definitions of which RBD sites are mutated in each variant are shown in Table 1.

**Table 1.** Mutated RBD sites in SARS-CoV-2 variants.

| variant | mutated RBD sites |
| --- | --- |
| Wuhan-Hu-1 |  |
| Alpha | 501 |
| Beta | 417 484 501 |
| Gamma | 417 484 501 |
| Delta | 452 478 |
| Lambda | 452 490 |
| Mu | 346 484 501 |
| Omicron | 339 371 373 375 417 440 446 477 478 484 493 496 498 501 505 |
| PMS20 | 346 417 440 445 455 475 484 501 |

## Acknowledgments

We thank collaborators in the Crowe lab, Nussenzweig lab, Bjorkman lab, and at Vir Biotechnology for sharing the antibodies that were used in prior published studies to generate the deep mutational scanning data aggregated here. We thank the Moyo and de Oliveira labs and other researchers in South Africa and Botswana for rapidly sharing information about the Omicron variant to enable the analyses in the last section of the results. This work was supported in part by the NIH/NIAID under R01AI141707 (to JDB) and T32AI083203 (to AJG). The work was also supported by the Gates Foundation grant INV-004949 to JDB. TNS is a Howard Hughes Medical Institute Fellow of the Damon Runyon Cancer Research Foundation. JDB is an Investigator of the Howard Hughes Medical Institute.

## Competing interests

JDB consults for Moderna, Flagship Labs 77 and Oncorus. JDB is an inventor on a Fred Hutch licensed patents related to deep mutational scanning of viral proteins.

## Notes

https://jbloomlab.github.io/SARS2_RBD_Ab_escape_maps/escape-calc/

https://github.com/jbloomlab/SARS2_RBD_Ab_escape_maps

https://jbloomlab.github.io/RBD_escape_calculator_paper/

